# Pathogenicity Patterns in Cytochrome P450 Family

**DOI:** 10.1101/2025.03.30.646180

**Authors:** Anna Špačková, Nina Kadášová, Ivana Hutařová Vařeková, Karel Berka

## Abstract

**Motivation:** Cytochrome P450 proteins play a crucial role in human metabolism, from the production of hormones to drug metabolism. While multiple commonly known variants have known effects on the individual cytochrome P450 protein performance, the pathogenicity information is usually experimentally limited to only a few mutations. Current pathogenicity prediction software allows one to extend the scope to virtually mutate all amino acids with missense mutations. In this work, we do a comprehensive exploration that unveils pathogenicity patterns in the human cytochrome P450 family. Pathogenicity analysis was conducted across proteins using SIFT and AlphaMissense algorithms.

**Results:** Our findings indicate a progressive increase in pathogenicity along protein tunnels-identified via MOLE-toward the cofactor binding site, underscoring the essential role of cofactor interactions in enzymatic function. Notably, tunnel integrity emerges as a critical factor, with even single amino acid alterations potentially disrupting molecular guidance to active sites. These insights highlight the fundamental role of structural pathways in preserving cytochrome P450 functionality, with implications for understanding disease-associated variants and drug metabolism.

**Availability:** Data and source code can be found at https://github.com/annaspac/P450_pathogenicity_codes

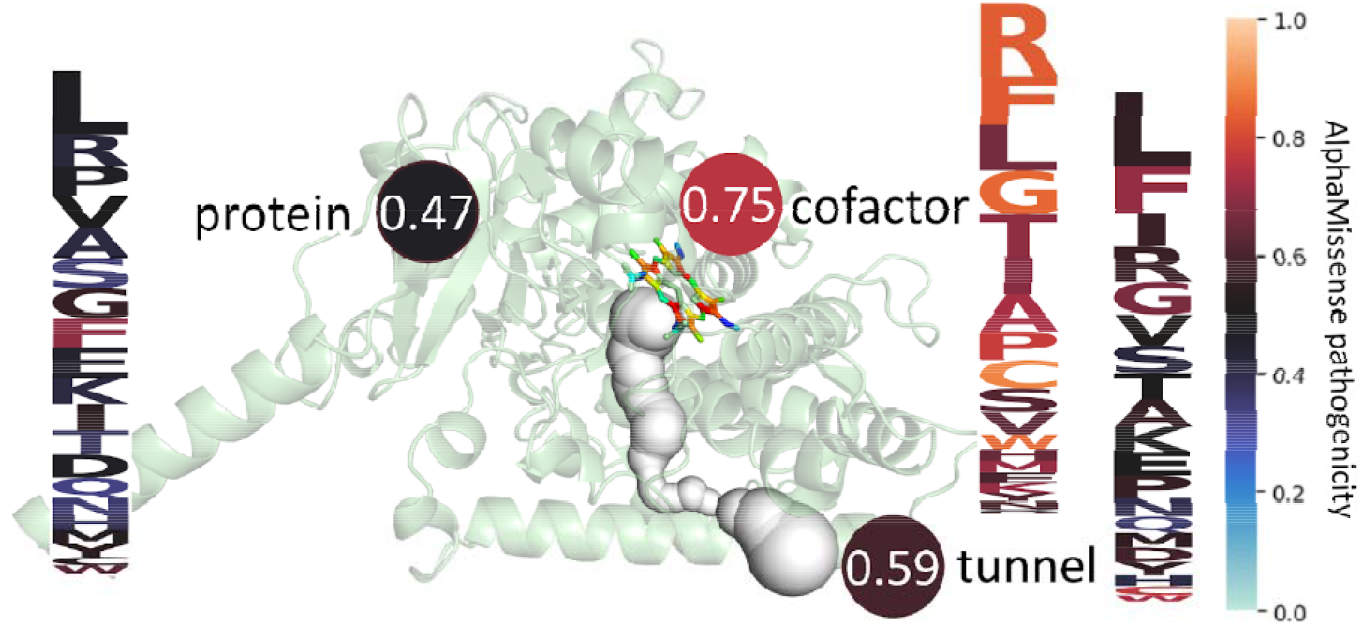

## 1 Introduction

The human cytochrome P450 (CYP) family comprises 57 genes, with diverse roles in drug metabolism, processing of foreign chemicals, synthesis and metabolism of vitamin D3 (Sawada *et al*. 1999, Kitanaka *et al*. 1998), calcium and phosphorus homeostasis (Tang *et al*. 2012), **re**. It emphasizes their pivotal role in maintaining physiological integr**ity**. (Nebert *et al*. 2013)

Tunnels within biomacromolecular structures, encompassing proteins, nucleic acids, and their complexes, serve crucial biological functions by facilitating the movement between internal spaces and the external environment. (Huang *et al*. 2001) Within enzymes, these tunnels play an important role in permitting the transit of substrates to and from buried active sites, contributing to essential biochemical processes. (Pravda *et al*. 2014) In essence, the presence of a tunnel streamlines the selection of substances that are allowed to enter specific regions within the protein amidst the complex mix of molecules in the cell. (Marques *et al*. 2017) Tunnels represent significant structural features that may contribute to the control of enzymatic functions and other biological processes. (Ljungdahl *et al*. 2023) Cofactors serve as redox carriers in biosynthetic and catabolic reactions, crucial in transferring energy within the cell. (Strushkevich *et al*. 2010) The majority of cofactors are non-covalently attached to proteins. There are, however, some proteins in which the cofactor binds covalently. (Stevens *et al*. 2005)

Genome sequencing has unveiled significant genetic diversity in human populations. (Karczewski *et al*. 2020) Mutations can modify protein amino acid sequences and structure. Pathogenic ones disrupt protein function, while benign variants have minimal effects. (Wang *et al*. 2013) Therefore, in this study, we focused on the pathogenicity of entire proteins, amino acids around tunnels, and cofactors. Our prediction is that any amino acid substitution near tunnels and cofactors should be more pathogenic, thereby having a greater impact on the overall protein function.

## 2 Methods

### 2.1 Protein structure file and compute tunnels

For the identification of P450 cytochromes in the UniProt (UniProt Corsortium 2023) database, the results encompass 60 such proteins. Their IDs were recorded and searched in the AlphaFill (Hekkelman *et al*. 2023) database, where they stored AI-folded protein structures enriched with cofactors. Protein tunnels were quantified using the desktop MOLE (Sehnal *et al*. 2013) algorithm (web MOLE could also be used (Pravda *et al*. 2018)) on these structures, providing insights into the amino acid composition around the tunnels and the physicochemical properties of these pathways. Interestingly, according to the algorithm, two proteins in the dataset did not exhibit any tunnels.

### 2.2 Pathogenicity of amino acids

The AlphaMissense (Cheng *et al*. 2023) algorithm computed pathogenicity values for the dataset. Subsequently, the SIFT (Ng and Henikoff 2003) algorithm was applied, and the results for twenty possible amino acid substitutions were averaged for each residue. The average pathogenicity values were then calculated for each protein using both the SIFT and AlphaMissense methods.

Further analysis focused on amino acids near tunnels identified by the MOLE algorithm, with their respective pathogenicity values determined. Similarly, for amino acids around cofactors, the nearest amino acids were identified for each cofactor atom, and the average pathogenicity was computed from the twenty representatives. This comprehensive approach provided insights into the average pathogenicity for entire proteins, the vicinity of tunnels, and the regions around cofactors.

All results were processed and obtained using codes programmed in Python 3.9.13, utilizing libraries such as os, json, pandas, seaborn, and matplotlib.pyplot.

## 3 Results

After categorizing pathogenicity based on individual residues, there is an observed increased occurrence of amino acids with higher pathogenicity values for residues found around tunnels and cofactors. The occurrence of values representing benign mutations is significantly suppressed in protein-tunnel and tunnel-cofactor scenarios. This holds true for the distribution of amino acids in both AlphaMissense and SIFT calculations **Figure 1**.

**Fig. 1.**
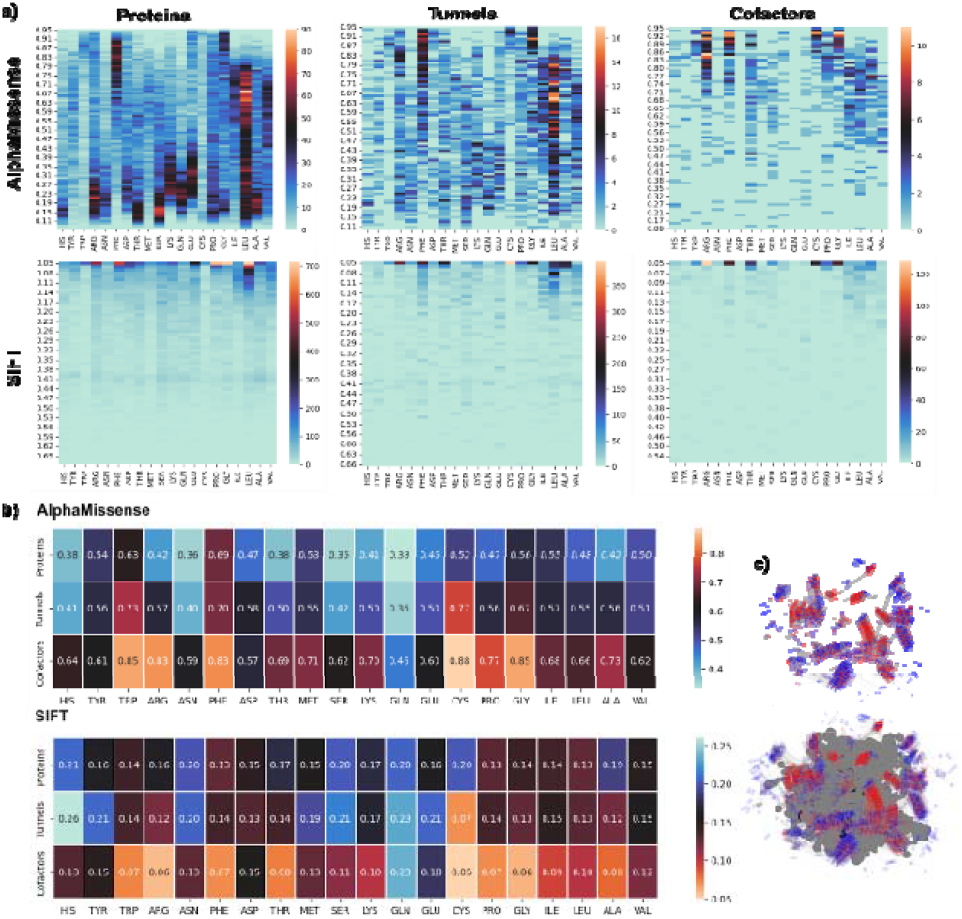
a) Pathogenicity in human cytochrome P450, around tunnels, and cofactors assessed using AlphaMissense and SIFT, heatmap illustrates the frequency of occurrence of specific amino acids with their corresponding pathogenicity values. AlphaMissense results show a range of pathogenicity for entire proteins, from 0 (benign) to 1 (pathogenic), with residues around tunnels mostly at higher values. This pattern intensifies around cofactors. The SIFT algorithm maintains this trend, but most amino acids are designated as pathogenic, particularly around the value of 0.05 in this case, from 0 (pathogenic) to 1 (benign). b) Comparison of average pathogenicity in entire proteins, around tunnels, and near cofactors in cytochromes P450. In proteins, phenylalanine and tryptophan exhibit the highest pathogenicity, followed by glycine. Around tunnels, the trend changes, with cysteine showing the highest pathogenicity, followed by tryptophan and phenylalanine with consistently high pathogenicity. Near cofactors, cysteine, glycine, tryptophan, arginine, and phenylalanine are the most pathogenic. Overall, the trend persists, with average pathogenicity increasing from proteins, through tunnels, to cofactors, for each represented amino acid. c) Overlaid structures of cytochrome P450 proteins in 2DProts displaying their pathogenicity levels according to the AlphaMissense algorithm. Redder areas indicate regions with higher pathogenicity, potentially indicating clusters of pathogenic mutations within specific regions of the proteins. In the structure, tunnels are depicted across all proteins.

The scale of pathogenicity differs between SIFT and AlphaMissense algorithms. AlphaMissense ranges from zero, representing benign substitutions, to one, representing pathogenic substitutions. Values observed in P450 cytochrome proteins are distributed more or less regularly. However, patterns emerge showing an increased occurrence of amino acids with certain pathogenicity. Elevated occurrences above a value of 0.5, combined with a large number of such pathogenic amino acids, are observed for phenylalanine, glycine, isoleucine, leucine, and valine across the entire protein. In tunnels, the occurrence is suppressed at more benign values, and around cofactors, these amino acids are nearly absent.

Conversely, in the SIFT algorithm, the scale is inverted, with most amino acids being designated as highly pathogenic. However, a reduced presence of less pathogenic amino acids is observed around tunnels and, to a greater extent, around cofactors, where they are almost absent **Figure 1**. In entire protein structures, amino acids with the highest pathogenicity, such as leucine, glycine, proline, phenylalanine, and tryptophan, are most abundant. Notably, around cofactors, glycine predominates in pathogenicity, followed by arginine, phenylalanine, and threonine. Additionally, arginine, phenylalanine, glycine, cysteine, threonine, and proline are prominently represented among cofactors. Alanine, leucine, and isoleucine are also present, albeit to a lesser extent.

A comparison of the average pathogenicity of individual amino acids was conducted. The findings from AlphaMissense conform to the protein-tunnel and tunnel-cofactor patterns, with one exception: aspartic acid, where the difference is only 0.01. Among the most pathogenic amino acids in the entire protein were phenylalanine, tryptophan, glycine, isoleucine, and tyrosine. Around tunnels, the order of the most pathogenic amino acid averages was cysteine, tryptophan, phenylalanine, and glycine. Amino acids in the vicinity of cofactors, including cysteine, tryptophan, glycine, phenylalanine, and arginine, exhibited some of the highest levels of pathogenicity. Overall, if an amino acid ranks among the most pathogenic in one of the groups- whole protein, tunnel vicinity, or cofactor vicinity-it tends to display higher pathogenicity in the other groups as well.

The SIFT results maintain the pattern for 11 amino acids. Among the amino acids with the highest average pathogenicity in whole proteins are leucine, proline, phenylalanine, isoleucine, glycine, and tryptophan. In the tunnel region, these are followed by cysteine, arginine, serine, aspartic acid, leucine, and glycine. Within the cofactor vicinity, the most pathogenic amino acids are cysteine, glycine, phenylalanine, tryptophan, and proline **Figure 1**.

Both tools agree on the amino acids phenylalanine, isoleucine, glycine, and tryptophan in whole proteins, cysteine and glycine around tunnels, and cysteine, glycine, phenylalanine, and tryptophan around cofactors **Figure 1**. The number of individual amino acids around the tunnels varied significantly, with leucine and phenylalanine being the most abundant, comprising around 15%. Approximately 7% representation was observed for isoleucine and arginine, while valine accounted for 6%. The least represented amino acids around the tunnels were aspartic acid and histidine.

AlphaMissense adheres to the average pathogenicity pattern in the protein, tunnels, and cofactors environment at 93.1%, while SIFT determination adheres to the pattern only at 67.3%. For more accurate results, predictions can be made using the AlphaMissense algorithm, which is among the most effective pathogenicity prediction algorithms. The average pathogenicity value for proteins calculated using SIFT ranges from 0.05 to 0.26, while for AlphaMissense, it spans from 0.29 to 0.91. The wider range used with AlphaMissense ensures a more accurate determination of pathogenicity. Examining the phylogenetic tree, the average pathogenicity value correlates with relatedness in the phylogenetic tree, especially among proteins within the same branch **Figure 2**.

**Fig. 2.**
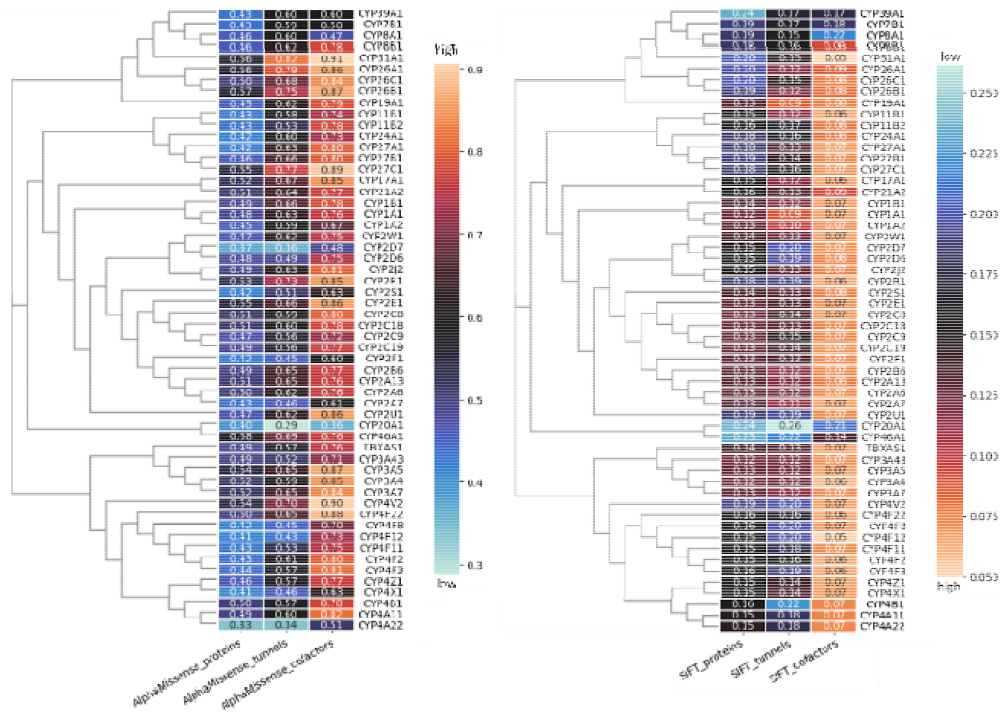
Comparing the average pathogenicity in P450 cytochromes, focusing on entire proteins, tunnels, and cofactor surroundings, with protein phylogenetics. Most of the protein follows the pattern of increasing average pathogenicity from the entire protein through tunnels to cofactors. For AlphaMissense results, four do not follow the pattern of increasing pathogenicity from protein, through tunnels, to cofactors. Two do not exhibit increased pathogenicity around tunnels compared to the entire protein, and in one case, there is no higher pathogenicity for the entire protein compared to the cofactor surroundings. The average pathogenicity for all but two is higher around tunnels compared to the entire protein. With two exceptions, the average pathogenicity increases around cofactors compared to the average pathogenicity in tunnels. With SIFT, the preservation of pathogenicity is clearly visible in individual branches of relatedness for the average pathogenicity of the protein.

The correlation of protein pathogenicity, tunnels, and cofactors is particularly evident in the results from SIFT, which stems from the algorithm’s reliance on related sequences. It’s fascinating to observe that individual branches adhere to pathogenicity within certain ranges, notably in learning the pathogenicity of the entire protein, where a similarity in pathogenicity is preserved within the same branches of the phylogenetic tree. Near cofactors, this value remains largely unchanged except for two branches, which differ significantly **Figure 2**.

In AlphaMissense, there is a certain similarity in the values of pathogenicity, but also occasional exceptions, which typically consist of three distinct values in the columns representing pathogenicity for the entire protein, tunnels, and around cofactors.

## Discussion

Cytochromes P450 are essential for the proper functioning of the human body, appearing in several physiological processes associated with various diseases. The calculation of protein tunnels using the MOLE tool provides information about the physicochemical properties of tunnels, which subsequently influences the types of molecules that can enter the tunnels and react with the buried active site. Studying the pathogenicity of different parts of cytochromes provides information about their importance. The study examines the pathogenicity of individual amino acids present in proteins. One of the key findings in this study is the increasing pathogenicity in protein-tunnel and tunnel-cofactor pairs, this increase suggests that amino acid substitutions across the entire protein are not as severe as those occurring near tunnels and even more so in the vicinity of cofactors. This substitution can affect the physicochemical properties at the sites where transport and reactions with the active site occur in the protein.

The comparison of results from the AlphaMissense and SIFT algorithms provides information on the reliability and accuracy of pathogenicity prediction outcomes. While both algorithms agree on certain amino acids with high pathogenicity, such as phenylalanine, glycine, cysteine, and tryptophan, in the entire protein as well as in the regions of tunnels and cofactors, there are noticeable differences. Pathogenicity values range from 0 to 1, with AlphaMissense designating 1 as pathogenic and 0 as benign, whereas with the SIFT algorithm, it’s the opposite. Results are distributed across the full scale in AlphaMissense, but with SIFT, pathogenicity for most amino acids is determined up to a value of 0.5.

The phylogenetic tree within the cytochrome P450 family shows a certain correlation with pathogenicity patterns in the case of related protein sequences. Preservation of pathogenicity in specific branches of the tree is particularly evident in the results of the SIFT algorithm, which may be attributed to how the algorithm determines pathogenicity and its close association with multiple sequence alignment of proteins (Sim *et al*. 2012). Overall, the preservation of pathogenicity in branches underscores the functional significance of related sequences.

The analysis of amino acid occurrence around tunnels and cofactors reveals an increased presence of pathogenicity and specific amino acids, further emphasizing the importance of tunnels for transporting small molecules to the active site.

## Acknowledgements and Funding

This work has been supported by the Palacký University Olomouc [IGA_PrF_2025_003 to A.Š., N.K.], IT4INNOVATIONS [OPEN-28-49 to A. Š.]. Computational resources were also provided by the e-INFRA CZ project, supported by the Ministry of Education, Youth and Sports of the Czech Republic.

